# Predicting Lung Cancer in Korean Never-Smokers with Polygenic Risk Scores

**DOI:** 10.1101/2022.11.23.515119

**Authors:** Juyeon Kim, Young Sik Park, Jin Hee Kim, Yun-Chul Hong, Young-Chul Kim, In-Jae Oh, Sun Ha Jee, Myung-Ju Ahn, Jong-Won Kim, Jae-Joon Yim, Sungho Won

## Abstract

In the last few decades, genome-wide association studies (GWAS) with more than 10,000 subjects have identified several loci associated with lung cancer. Hence, recently, genetic data have been used to develop novel risk prediction tools for cancer. The present study aimed to establish a lung cancer prediction model for Korean never-smokers using polygenic risk scores (PRSs). PRSs were calculated using a thresholding-pruning-based approach based on 11 genome-wide significant single nucleotide polymorphisms (SNPs). Overall, the odds ratios tended to increase as PRSs were larger, with the odds ratio of the top 5% PRSs being 1.71 (95% confidence interval: 1.31−2.23), and the area under the curve (AUC) of the prediction model being of 0.76 (95% confidence interval: 0.747−0.774). The receiver operating characteristic (ROC) curves of the prediction model with and without PRSs as covariates were compared using DeLong’s test, and a significant difference was observed. Our results suggest that PRSs can be valuable tools for predicting the risk of lung cancer.

## 1 INTRODUCTION

Lung neoplasms are the leading cause of cancer worldwide (Fitzmaurice et al., 2019), with lung cancer having been the second most commonly diagnosed cancer in 2020, after breast cancer in women (Bray et al., 2018). In Korea, the age-adjusted prevalence of lung cancer in 2018 was 94.1 cases per 100,000 people, accounting for 4.7% of all cancer cases, and its age-adjusted incidence rate was of 28.0 cases per 100,000 people, accounting for 11.7% of all cancers (Korea Central Cancer Registry, 2020). Among men, the incidence of lung cancer in 2018 was 41.9 cases per 100,000 people, which is the second highest in Korea, whereas in Japan, United States, and United Kingdom the mean incidence was of 41.4, 40.1, and 35.5 cases per 100,000 people, respectively (Korea Central Cancer Registry, 2020). Lung cancer is the most common cancer affecting men aged > 65 years in Korea (Korea Central Cancer Registry, 2020), despite the proportion of never-smokers increased to 25.4% in 2009-2012, which was 19.1% in 2004-2008 (Park & Jang, 2016)

Smoking is a major risk factor for the progression of lung cancer and has been associated with over 80% of lung cancer cases in the Western world (Corrales et al., 2020). Indeed, reduced smoking habits has led to a decrease in mortality and incidence of lung cancer (Thandra et al., 2021). Nonetheless, even though most patients are smokers, the proportion of never-smokers with lung cancer has been increasing over time (Couraud et al., 2012), with the World Health Organization estimating that 25% of lung cancer cases worldwide occur in never-smokers (Ferlay et al., 2010). Therefore, the risk profiles of never-smokers are expected to be markedly different from those of smokers, with family history, secondhand smoke, cooking oil fumes, radon exposure, domestic fuel smoke, asbestos, and menopausal hormone replacement therapy being suggested as potential risk factors associated with lung cancer in never-smokers. However, to date, none of these suggested risk factors have been exclusively identified in never-smokers (Couraud et al., 2012; Hung et al., 2021).

Lung cancer in never-smokers has been considered to be a distinct medical entity from that in ever-smokers, and some clinically important features have been identified. First, it is more frequent in certain regions than in others (Asia > North America > Europe) (Couraud et al., 2012). Second, mutations in the epidermal growth factor receptor (EGFR) are more common in 1) adenocarcinomas than in non-small cell lung cancer and 2) in never-smokers than in ever-smokers (Chapman et al., 2016). Noteworthily, although the somatic variant profile of Asian populations is very similar to that of Europeans, the prevalence of *EGFR* mutations is higher in Asian women than in Caucasian women (Chapman et al., 2016). Therefore, it is unlikely that other risk factors, such as secondhand smoke, could be responsible for the increased lung cancer incidence in the Asian population (Mitsudomi, 2014). Several studies have suggested that *EGFR* mutations occur independently of smoking and that the low frequency of *EGFR* mutations in smokers could be explained by the occurrence of smoking-related lung cancer (Mitsudomi, 2014; Shi et al., 2014; Truong et al., 2010). Moreover, genome-wide data related to *EGFR* mutations failed to provide relevant knowledge on carcinogenesis.

GWAS have discovered many common genetic variants associated with complex traits and disorders (Buniello et al., 2019; Klein et al., 2005; Visscher et al., 2017). Most cancers are highly polygenic (Stahl et al., 2012; Zeng et al., 2018; Zhang et al., 2018; Zhang et al., 2020), with lung cancer being one of the most polygenic (Zhang et al., 2020), along with breast and oropharynx cancers. For these polygenic traits, the effect size associated with each risk variant is small, and individuals with multiple risk variants tend to have an elevated disease risk (Chatterjee et al., 2013). Therefore, PRS can be useful for risk assessment as it combines multiple variants into scores that evaluate genetic susceptibility (Dudbridge, 2013). Several studies have shown that PRS can be used as a predictor of lung cancer in a population that includes both ever-smokers and never-smokers; however, most of these studies were conducted on non-Hispanic whites, and no study evaluated exclusively for Asian never-smokers. In the present study, a lung cancer prediction model for Korean never-smokers was built using PRSs and its accuracy was evaluated.

## 2 MATERIALS AND METHODS

### 2.1 Korean lung cancer cohorts

Never-smoking Korean lung cancer subjects were recruited from five different institutes: Seoul National University Hospital (SNUH), Yeonsei University (YSU), Sejong University (SU), Samsung Medical Center (SMC), and Chonnam National University (CNU) (Ahn et al., 2012; Kim et al., 2013; Lan et al., 2012; Lee et al., 2017). Data from non-smoking controls were obtained from the CAVAS study of the Korean Genome and Epidemiology Study (Kim & Han, 2017). Never-smokers were defined as those who had smoked less than 100 cigarettes in their lifetime or had never smoked. A total of 8,348 individual were included in the study (1,642 cases and 6,706 controls matched according to their principal component (PC) scores calculated using the EIGENSTRAT method (Price et al., 2006)). All participants provided written informed consent, and the study was approved by the institutional review board and ethics committee of the Seoul National University Hospital (approval no. H-1906-126-1042).

### 2.2 Genotyping, quality control, and imputation

SNUH and YSU cohorts were genotyped using Axiom KoreanChip V1.0 or Axiom KoreanChip V1.1 (Moon et al., 2019), SU and SMC cohorts were evaluated using the Affymetrix Genome-Wide Human SNP Array (5.0 and 6.0, respectively), and CNU cohort was evaluated using an Illumina Human660W-Quad array. Variant calling for SNUH and YSU was performed using the K-medoid approach (Seo et al., 2019). The analysis approach used is summarized in **Figure 1**. As different genotyping platforms can generate substantial numbers of false-positive data, and quality controls (QC) were carefully performed. SNPs were removed if call rates were < 95% or 99%,*P*-values for Hardy–Weinberg equilibrium were < 10^-3^ or 10^-5^, or minor allele frequencies (MAFs) were < 5%. Subjects were also excluded if there was sex inconsistency, call rate < 0.95, outlying heterozygosity (heterozygosity rate > mean ± 3 standard deviation [SD]), or estimated identity-by-descent > 0.9. QCs were conducted with cases and controls separately for each participant institution, with cases and controls from the same institute being merged and the same QC being applied to the merged data with the following additional step: SNPs were removed if missing rates between cases and controls differed significantly (*P* < 0.01). Lastly, genotyping platforms of SNUH and YSU, and for SU and SMC were the same, and subjects from institutes with the same genotyping platforms were pooled. The same QCs were applied to pooled subjects. After QCs, the remaining subjects and SNPs were used to impute the untyped SNPs using the Michigan imputation server. Non-Europeans of the Haplotype Reference Consortium (r1.1 2016) were selected as reference panel, and Eagle (v2.4) was used as the phasing program. Imputed SNPs were removed if MAFs were < 0.05, *R^2^* < 0.3, *P*-values for the Hardy–Weinberg equilibrium exact test were < 10^-3^ or 10^-5^, call rates were < 95% or 99%.

**Figure 1.**
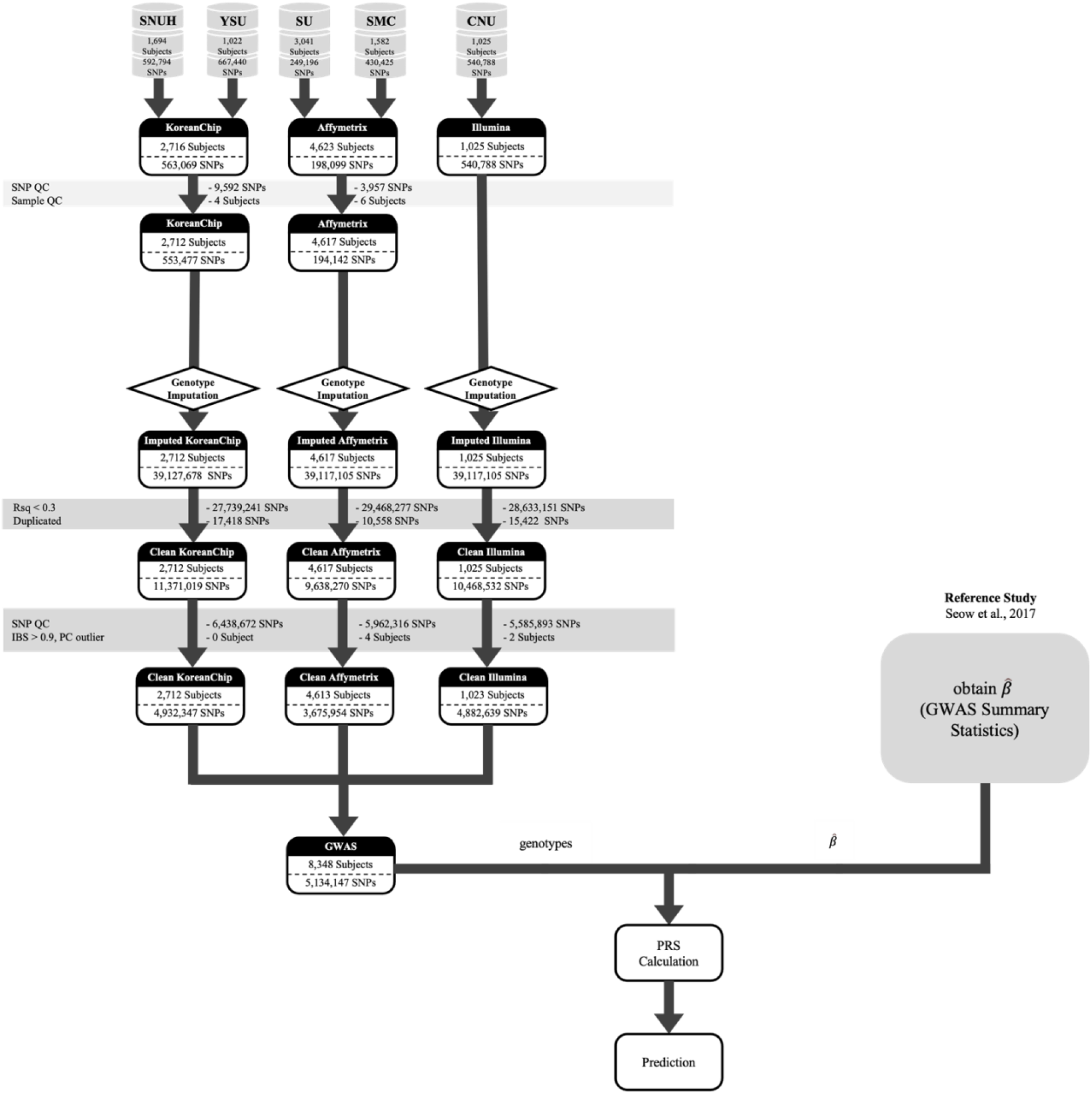
Flowchart of data collection and analysis protocols. Abbreviations: CNU, Chonnam National University; GWAS, genome-wide association study; IBS, identity by state; PC, principal component; PRS, polygenic risk score; QC, quality control; Rsq, R squared; SNP, single nucleotide polymorphism; SMC, Samsung Medical Center; SNUH, Seoul National University Hospital; SU, Sejong University; YSU, Yeonsei University.

Association analyses were conducted using logistic regression. To adjust for population stratification, PC scores were calculated, and the 10 PC scores corresponding to the 10 largest eigenvalues were included as covariates in the following logistic regressions:

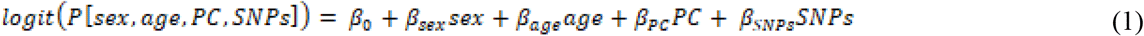

The genomic inflation factor and quantile-quantile plot were used to compare the genome-wide distribution of the test statistic for *H_0_: β_SNPs_ = 0* with the expected null distribution.

### 2.3 Polygenic risk score construction

Calculation of PRS requires an effect size estimate of genome-wide significant SNPs. GWAS Catalog (https://www.ebi.ac.uk/gwas/) and PubMed (https://pubmed.ncbi.nlm.nih.gov) were screened to obtain GWAS summary statistics of lung cancer in never-smokers of Asian ancestry. Seow *et al*. (Seow et al., 2017) conducted a GWAS using the largest East Asian population, reporting 11 genome-wide significant SNPs, among which the genotypes of 10 SNPs were available in each Korean cohort(Table S1); thus, their summary statistics were incorporated to build the PRS. Let 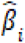 be the log odds ratios (ORs) obtained from Seow *et al*. (Seow et al., 2017) for SNP *i* (*i* = 1, […], 11) and *x_ij_* be the number of risk alleles of SNP *i* for subject *j* in the Korean cohort (*x_ij_* = 0, 1, 2); then, the PRS of subject *j* was calculated by a weighted sum of the risk alleles that an individual carries, as follows:

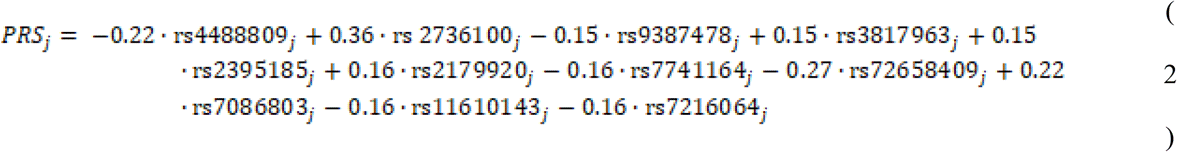

### 2.4 Logistic regression and its prediction accuracy

The prediction model was built with using logistic regression models. Lung cancer status was used as the response variable. PRSs were categorized into nine different groups based on PRSs of the subject percentiles of the controls: < 5%, 5–10%, 10–20%, 20–40%, 40–60%, 60–80%, 80–90%, 90–95%, and > 95%, which were indicated as 1, 2, (…), and 9, respectively. PRSs were incorporated as covariates to estimate their ORs after adjusting for sex, age, 10 PC scores, and genotyping array as follows:

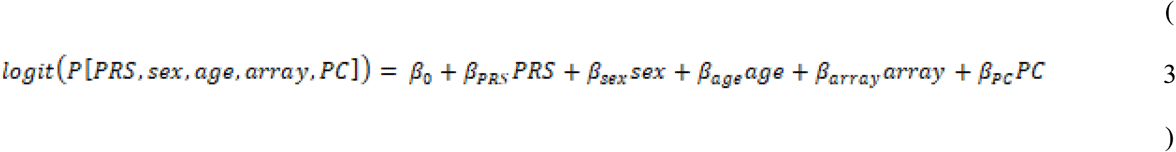

In this study, 10 PC scores were used to adjust the population stratification. To assess the ability of the PRS to identify high-risk cases, we considered an alternative model in which the PRS was coded as 1 for the top 1% PRSs, otherwise was coded as 0.

The prediction accuracy of the logistic regression was evaluated using the AUC. The confidence interval and *P*-value were obtained using the DeLong’s test. Subjects from SU and CNU overlapped with those of Seow *et al*. (Seow et al., 2017), and most never-smokers were females. Therefore, the accuracy of the prediction model was evaluated according to three different scenarios: Dataset 1, all subjects (SNUH, YSU, SU, SMC, and CNU); Dataset 2, only females; and dataset 3, subjects from SNUH, YSU, and SMC. Moreover, ROC curves of the prediction model with and without PRSs as covariates were compared using DeLong’s test. The ORs of the PRSs were estimated and adjusted for the first 10 PC scores, sex, age, and dataset. All the analyses were performed using Plink (v1.9 and v2.0), ONETOOL (Song et al., 2018), R (v3.6.3), and Python (v2.7.17).

### 2.5 Meta analyses

Meta-analyses were conducted to calculate the combined effect sizes for each SNP using METAL (Willer et al., 2010). The effect sizes of each SNP were combined using weighted means. Forest plots were obtained using R (v3.6.3)(Figure S1).

## 3 RESULTS

### 3.1 Descriptive statistics

The descriptive statistics of the subjects included in the study are shown in **Table 1**. Among the 8,348 individuals evaluated, 72.4% were females and a total of 84.6% of the patients were pathologically diagnosed with adenocarcinoma. Significant differences in age were observed among the different study cohorts.

**Table 1.** Descriptive statistics.

|  | SNUH | YSU | SU | SMC | CNU | Total | *P*-value |
| --- | --- | --- | --- | --- | --- | --- | --- |
| **Overall, *N*** | 1,694 | 1,018 | 3,038 | 1,575 | 1,023 | 8,348 |  |
| Case | 206 | 172 | 276 | 407 | 581 | 1,642 | < 0.001 |
| Control | 1,488 | 846 | 2,762 | 1,168 | 442 | 6,706 |  |
| **Sex, *N*** |  |  |  |  |  |  |  |
| Male | 389 | 336 | 318 | 339 | 0 | 1,382 | < 0.001 |
| Female | 1,305 | 682 | 2,720 | 1,236 | 1,023 | 6,966 |  |
| **Mean age (SD), years** | 53.5 (10.0) | 47.2 (11.6) | 53.3 (9.4) | 60.0 (8.1) | 60.5 (10.7) |  | < 0.001 |
| **Histology, *N*** |  |  |  |  |  |  |  |
| AD | 192 | NA | 240 | 359 | 453 |  | < 0.001 |
| Non-AD | 14 | NA | 36 | 48 | 128 |  |  |

Abbreviations: AD, adenocarcinoma; CNU, Chonnam National University; NA, not available; SD, standard deviation; SMC, Samsung Medical Center; SNUH, Seoul National University Hospital; SU, Sejong University; YSU, Yeonsei University.

### 3.2 Odds ratios of PRS and prediction accuracy

The PRSs were calculated for each dataset. The Kolmogorov-Smirnov test showed that the PRSs were normally distributed, and that the cases had significantly higher PRSs than controls in all datasets (**Table 2**, **Figure S2**). Moreover, Table 2 also shows that cases always have significantly higher means than controls in Dataset1, which includes all subjects (*P* = 4.50 × 10^-10^ for KoreanChip; *P* = 6.34 × 10^-9^ for Affymetrix; *P* = 3.63 × 10^-8^ for Illumina array). the estimated ORs of the PRSs tended to increase in higher percentile groups of the PRSs, compared with the reference percentile group (40–60%) (**Figure 2**). OR of the top 5% PRS group was 1.71 (95% confidence interval [CI]: 1.31–2.23; *P* = 7.40 × 10^-5^), and for females the OR was 1.66 (1.25–2.19; *P* = 3.76 × 10^-4^). The OR of the top 5% PRS group was maximized for Dataset 3 (OR = 2.45 [1.74–3.44]; *P* = 2.21 × 10^-7^) (**Table S2**). No significant differences between men and women were observed in Dataset1, which includes all subjects. The ORs of the bottom 5% PRS group were less than 1, indicating their protective effect against lung cancer (**Table S2**). Some PRS percentile groups were not significantly different from the reference group, but an increasing tendency in ORs was observed for all datasets (**Table S2**).

**Figure 2.**
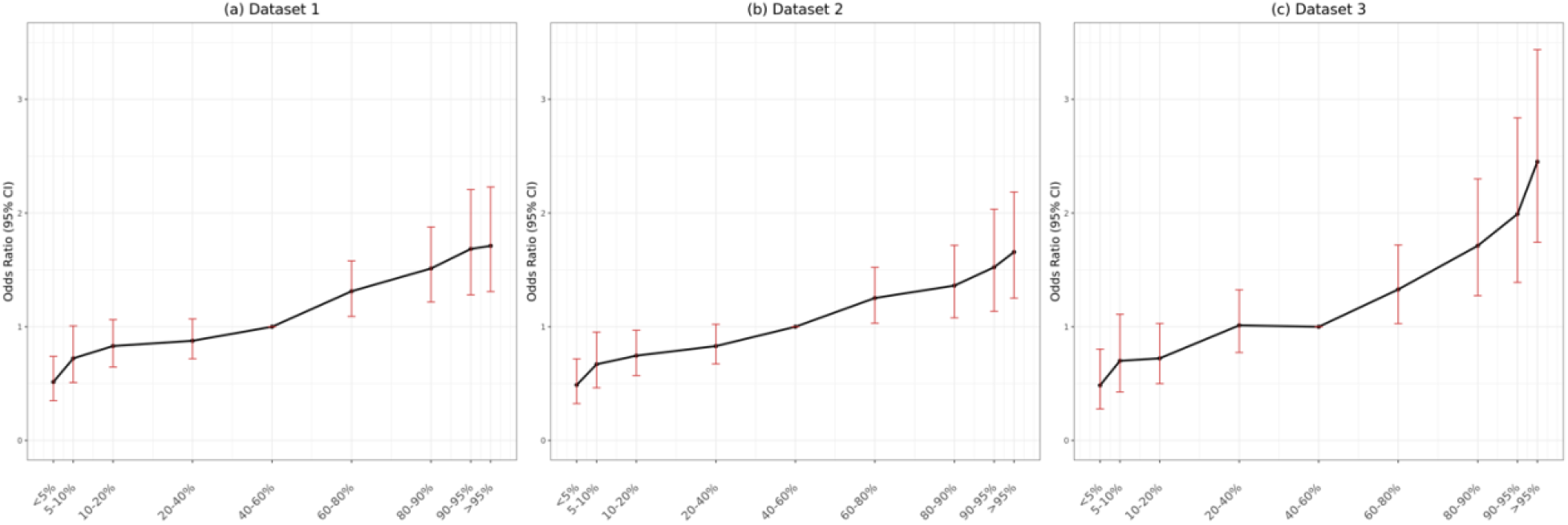
Odds ratios depending on percentiles of polygenic risk scores. Percentiles were defined in control subjects. Dots and vertical red lines represent the odds ratios and their 95% confidence intervals (CI), respectively. Middle quintile (40–60%) was considered as reference group. **(a)** Dataset 1 included all subjects from Chonnam National University (CNU), Samsung Medical Center (SMC), Seoul National University Hospital (SNUH), Sejong University (SU), and Yeonsei University (YSU). **(b)** Dataset 2 included only females from SNUH, YSU, SU, SMC, and CNU. **(c)** Dataset 3 included subjects from SNUH, YSU, and SMC.

**Table 2.** Mean differences between cases and controls

|  | Polygenic risk score, mean (SD) |  | KS test | *P*-values |  |
| --- | --- | --- | --- | --- | --- |
|  | Cases | Controls |  | *t*-test for phenotype | *t*-test for sex |
| Dataset 1<sup>†</sup> |  |  |  |  |  |
| KoreanChip | 0.30 (0.97) | −0.05 (1.0) | 0.33 | $4.50 \times 10^{-10}$ | 0.12 |
| Affymetrix | 0.20 (0.98) | −0.04 (1.0) | 0.08 | $6.34 \times 10^{-9}$ | 0.13 |
| Illumina | 0.15 (0.98) | −0.20 (0.99) | 0.66 | $3.63 \times 10^{-8}$ | NA<sup>†</sup> |
| Dataset 2<sup>‡</sup> |  |  |  |  |  |
| KoreanChip | 0.30 (0.95) | −0.04 (0.99) | 0.24 | $1.96 \times 10^{-8}$ | NA<sup>†</sup> |
| Affymetrix | 0.20 (0.98) | −0.05 (1.0) | 0.11 | $2.45 \times 10^{-8}$ | NA<sup>†</sup> |
| Illumina | 0.15 (0.98) | −0.20 (0.99) | 0.66 | $3.63 \times 10^{-8}$ | NA<sup>†</sup> |
| Dataset 3<sup>§</sup> |  |  |  |  |  |
| KoreanChip | 0.30 (0.97) | −0.05 (1.0) | 0.33 | $4.50 \times 10^{-10}$ | 0.12 |
| Affymetrix | 0.30 (1.04) | −0.10 (0.97) | 0.15 | $2.88 \times 10^{-10}$ | 0.80 |
| Illumina |  |  | NA |  |  |

<sup>†</sup>Included all subjects from Chonnam National University (CNU), Samsung Medical Center (SMC), Seoul National University Hospital (SNUH), Sejong University (SU), and Yeonsei University (YSU).
<sup>‡</sup>Included only females from SNUH, YSU, SU, SMC, and CNU.
<sup>§</sup>Included subjects from SNUH, YSU, and SMC.
Abbreviations: KS, Kolmogorov-Smirnov, NA, not available; SD, standard deviation.

Table 3 shows the ORs of the top 1% PRSs compared with the other PRS subgroups. Overall, the ORs were significant only for Dataset 3 and the OR of the top 1% PRS group was generally not higher than that of the top 5% group (**Table 3**, **Table S2**). For comparison, we referred to the ORs reported in the study by Fritsche *et al*. (Fritsche, 2020), which included data from lung cancer patients obtained from the UK biobank (UKB) and Michigan Genomics Initiative (MGI) (**Table 3**). Most of these patients were Caucasian, and both smokers and never-smokers were included. Although the ethnicity and smoking status were different, OR of the top 1% PRSs of Dataset3 was higher than those of UKB and MGI.

**Table 3.** Odds ratios of top 1%polygenic risk scores (PRSs) compared with the other PRS subgroups

|  | OR (95% CI) | *P*-value |
| --- | --- | --- |
| Dataset 1 | 1.29 (1.26–1.42) | 0.38 |
| Dataset 2 | 1.20 (1.26–1.43) | 0.53 |
| Dataset 3 | 2.22 (1.36–1.60) | 0.03 |
| UK Biobank | 1.75 (0.796–3.85) | NA |
| MGI | 1.94 (1.22–3.1) |  |

Abbreviations: CI, confidence interval; MGI, Michigan Genomics Initiative; NA, not available; OR, odds ratio.

### 3.3 Lung cancer prediction with polygenic risk scores

PRSs, age, and sex were considered covariates, and a prediction model was built. The pre dictors included sex, age, and continuous PRSs. The highest AUC was 0.764 (95% CI: 0.750–0.778; *P* = 7.02 × 10^-280^) from Dataset 2, followed by Dataset 1 (**Figure 3**, **Table 4**). AUCs from Dataset 1 and Dataset 2 were similar (*P* = 0.73), whereas those from Dataset 1 and Dataset 3 differed significantly (*P* = 1.80 × 10^-6^). Moreover, significant differences were observed concerning the ROC of the prediction model with and without PRSs as covariates in every scenario (**Table 4**).

**Figure 3.**
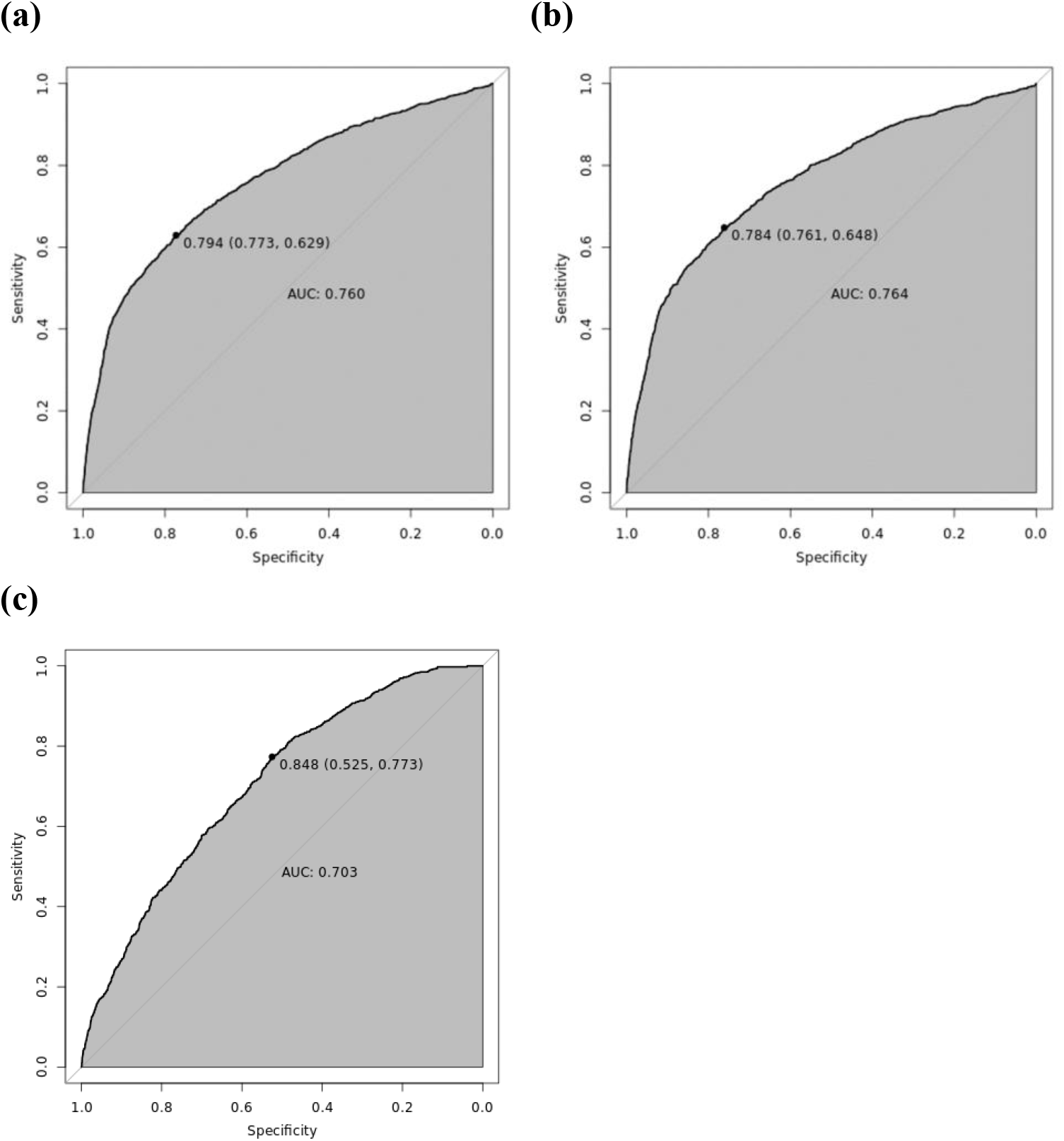
Receiver operating characteristic curves of the different datasets. **(a)** Dataset 1 included all subjects from Chonnam National University (CNU), Samsung Medical Center (SMC), Seoul National University Hospital (SNUH), Sejong University (SU), and Yeonsei University (YSU). **(b)** Dataset 2 included only females from SNUH, YSU, SU, SMC, and CNU. **(c)** Dataset 3 included subjects from SNUH, YSU, and SMC. AUC, area under the curve.

**Table 4.** Comparison of the area under the curves (AUC) of the prediction models with and without polygenic risk scores (PRSs) as a covariate

|  | AUC with PRSs<br>(95% confidence interval) | AUC without PRSs<br>(95% confidence interval) | *P*-value, DeLong test |
| --- | --- | --- | --- |
| Dataset 1 | 0.760 (0.747–0.774) | 0.750 (0.736–0.765) | $1.87 \times 10^{-5}$ |
| Dataset 2 | 0.764 (0.750–0.778) | 0.754 (0.739–0.769) | $4.01 \times 10^{-5}$ |
| Dataset 3 | 0.703 (0.684–0.722) | 0.673 (0.654–0.692) | $1.09 \times 10^{-7}$ |

## 4 DISCUSSION

Genetics for lung cancer in never-smokers has been considered one of the most important risk factors, with several studies having been performed to predict lung cancer while considering specific genetic factors. Recently, PRS has been applied to predict lung cancer; however, there no such studies have been conducted exclusively for Asian never-smokers. In this study, we constructed PRSs based on recent meta-analyses and evaluated their prediction accuracy for lung cancer in never-smokers in Korea. Our results show that individuals with PRSs higher than the reference percentile group(40-60%) have a much higher probability of developing lung cancer; thus, these PRSs can be considered valuable predictors of lung cancer.

To date, the largest studies exploring PRS in patients with lung cancer were based on the MGI and UKB cohorts, which consist of non-Hispanic white European populations, including ever and never-smokers. Their data suggested that the top 1% PRS represented an increased risk of lung cancer, with ORs of 1.75 for MGI (95% CI 0.796–3.85) and 1.94 for UKB (95% CI 1.22–3.1) groups (Fritsche, 2020). In our analyses, the OR of the top 1% PRS in never-smoker subjects of the Dataset 3 was 2.22 and was significantly associated with increase lung cancer risk (*P* = 0.03). For Datasets 1 and 2, the results were not statistically significant, but the data suggested a tendency for increased risk (OR > 1). These results indicate that the PRS can be utilized as a prognostic tool for lung cancer, regardless of the smoking status of the patient. Moreover, PRS can be useful for identifying never-smoking individuals with a high risk of lung cancer.

Our data showed that the predictive potential of PRS was similar between women and men. Multiple studies have shown substantial differences in lung cancer incidence according to sex. For example, studies based on data from The Cancer Genome Atlas showed that 15% of autosomal genes have sex-biased copy number alterations in several cancers, which can be associated with different mRNA expression profiles (Lopes-Ramos et al., 2020). Sex-related behavior and exposure can also affect gene mutations. In non-small cell lung cancer, the mutational spectrum of *EGFR* and *TP53* is influenced by sex. Indeed, the frequency of transversion mutations on *TP53* is 40% among women, which is higher than among men (25-28%) (Lopes-Ramos et al., 2020). Moreover, different methylation patterns by sex have been observed in various human tissues, such as blood, brain, and muscle (Lopes-Ramos et al., 2020). Therefore, it is expected that a combination of genetic mechanisms can contribute to epidemiological sex differences. However, in the present study, no sex-specific differences were observed. The largest difference in MAFs between men and women was 0.002, and there were no SNPs with minor allele frequencies that were significantly different between women and men.

This study had some limitations. First, although more than 1,500 lung cancer patients were considered, genetic analyses usually require more than 10,000 subjects, and the sample size may not be sufficient for evaluating the accuracy of the risk prediction model estimates using PRSs. Second, lung cancer consists of etiologically heterogeneous subtypes; however, this information was not available for this study. If etiological subtypes are to be considered, prediction accuracy may be greatly improved. Thus, further studies considering the subtype-specific genetic architecture of lung cancer with adequate sample size are still needed to further confirm our findings. Third, summary statistics were only available for some SNPs. Some studies showed that the AUC difference between the prediction model with 592,000 SNPs and 10 SNPs was 0.03, with inclusion of more SNPs not substantially improving prediction accuracy (Liyanarachchi et al., 2020). However, this conclusion was obtained based on non-Hispanic white populations; hence, further investi gations are necessary for East Asians. Fourth, it has been shown that secondhand smoke significantly affects lung cancer; however, its effect was not considered in this study. The mean age of the study participants was 60.2 years old. Laws to ban or stop smoking in all enclosed workplaces or cessation of health have been recently adopted, and the controls of our study participants may have also been affected by secondhand smoke.

Lung cancer has been widely known to be a highly heterogeneous disease that can occur and progress due to the interplay between permanent genetic mutations and epigenetic alterations (Dong et al., 2017). The lung cancer prediction accuracy can be improved by combining other clinical or lifestyle risk factors. In this study, we demonstrate that PRS can be a valuable tool for identifying individuals at a high risk of lung cancer. However, the predictive accuracy of PRSs is still not sufficiently good so it can be used in clinical practice; hence, further studies are warranted to improve it.

## DATA AVAILABILITY

The data that support the findings of this study are available from the corresponding author upon reasonable request.

## AUTHOR CONTRIBUTIONS

Juyeon Kim conceived the project, carried out data analysis, and drafted the manuscript. Sungho Won reviewed the literature and provided feedback. Young Sik Park, Jin Hee Kim, Yun-Chul Hong, Young-Chul Kim, In-Jae Oh, Sun Ha Jee, Myung-Ju Ahn, Jong-Won Kim, and Jae-Joon Yim conducted the epidemiological studies, contributed samples to the genotyping. All authors contributed to the writing of the manuscript.

## ACKNOWLEDGEMENTS

This work was supported by the Technology Innovation Program (20016417) funded by the Ministry of Trade, Industry, and Energy (Korea).

## CONFLICTS OF INTEREST

The authors declare no conflict of interest.

## Notes

### Competing Interest Statement

The authors have declared no competing interest.

