## Supplemental Table 1 as Table S1 for "Predicting Lung Cancer in Korean Never-Smokers with Polygenic Risk Scores"

**Supplementary Tables**

**Table S1.** Risk SNPs used for risk score assessment.

**Table S2.** Odds ratios by polygenic risk score percentile.

**Supplementary Figures**

**Figure S1.** Estimated odds ratios of the single nucleotide polymorphisms analyzed in each study.

**Figure S2.** Distributions of polygenic risk scores.

**Table S1.** Risk SNPs used for risk score assessment

| SNP ID† | Gene | Chr. | Physical Position | Major Allele | Risk Allele | Beta | OR | Subject | *P*-value |
|---|---|---|---|---|---|---|---|---|---|
| rs4488809 | *TP63* | 3 | 189,356,261 | T | C | −0.22314 | 0.8 | 14,455 | $4.30 \times 10^{-17}$ |
| rs2736100 | *TERT* | 5 | 1,286,516 | A | C | 0.357674 | 1.43 | 14,575 | $6.12 \times 10^{-43}$ |
| rs7741164 | *FOXP4* | 6 | 41,493,412 | G | A | 0.157004 | 1.17 | 21,179 | $3.96 \times 10^{-13}$ |
| rs9387478 | *ROS1/DCBLD1* | 6 | 117,786,180 | C | A | −0.15082 | 0.86 | 17,992 | $5.25 \times 10^{-11}$ |
| rs3817963 | *BTNL2* | 6 | 32,368,087 | T | C | 0.14842 | 1.16 | 14,000 | $1.63 \times 10^{-7}$ |
| rs2395185 | *HLA-DRB9* | 6 | 32,433,167 | G | T | 0.14842 | 1.16 | 17,394 | $2.04 \times 10^{-9}$ |
| rs2179920 | *HLA-DPB1* | 6 | 33,058,874 | C | T | 0.157004 | 1.17 | 14,477 | $1.69 \times 10^{-5}$ |
| rs72658409 | NA | 9 | 22,160,087 | C | T | −0.2744368 | 0.76 | 21,718 | $2.37 \times 10^{-10}$ |
| rs7086803 | *VTI1A* | 10 | 114,498,476 | G | A | 0.2231436 | 1.25 | 17,878 | $9.22 \times 10^{-17}$ |
| rs11610143 | NA | 12 | 52,349,071 | C | G | −0.1625189 | 0.85 | 20,901 | $3.55 \times 10^{-13}$ |
| rs7216064 | *BPTF* | 17 | 65,898,809 | A | G | −0.16252 | 0.86 | 16,350 | $6.19 \times 10^{-9}$ |

Abbreviations: Chr, chromosome; OR, odds ratio; SNP, single nucleotide polymorphism.

†Variants were selected from Seow *et al*. (*Hum Mol Genet.* 2017;26(2):454-65).

**Table S2.** Odds ratios by polygenic risk score percentile

| | Dataset 1† | | Dataset 2‡ | | Dataset 3§ | |
|---|---|---|---|---|---|---|
| | OR (95% CI) | *P*-value | OR (95% CI) | *P*-value | OR (95% CI) | *P*-value |
| < 5% | 0.51 (0.35–0.74) | $4.83 \times 10^{-4}$ | 0.49 (0.32–0.72) | $3.87 \times 10^{-4}$ | 0.48 (0.28–0.80) | $7.27 \times 10^{-3}$ |
| 5–10% | 0.72 (0.51–1.00) | $6.10 \times 10^{-2}$ | 0.67 (0.46–0.95) | $2.92 \times 10^{-2}$ | 0.70 (0.43–1.11) | $1.43 \times 10^{-1}$ |
| 10–20% | 0.83 (0.65–1.06) | $1.43 \times 10^{-1}$ | 0.75 (0.57–0.97) | $3.00 \times 10^{-2}$ | 0.72 (0.50–1.03) | $7.68 \times 10^{-2}$ |
| 20–40% | 0.88 (0.72–1.07) | $1.93 \times 10^{-1}$ | 0.83 (0.67–1.02) | $8.01 \times 10^{-2}$ | 1.01 (0.77–1.33) | $9.28 \times 10^{-1}$ |
| 40–60% | | | Reference | | | |
| 60–80% | 1.31 (1.09–1.58) | $4.00 \times 10^{-3}$ | 1.25 (1.03–1.52) | $2.33 \times 10^{-2}$ | 1.33 (1.03–1.72) | $3.02 \times 10^{-2}$ |
| 80–90% | 1.51 (1.22–1.89) | $1.80 \times 10^{-4}$ | 1.36 (1.08–1.72) | $9.10 \times 10^{-3}$ | 1.71 (1.26–2.30) | $3.57 \times 10^{-4}$ |
| 90–95% | 1.68 (1.28–2.20) | $1.72 \times 10^{-4}$ | 1.52 (1.14–2.03) | $4.60 \times 10^{-3}$ | 1.99 (1.39–2.84) | $1.50 \times 10^{-4}$ |
| > 95% | 1.71 (1.31–2.23) | $7.40 \times 10^{-5}$ | 1.66 (1.25–2.19) | $3.76 \times 10^{-4}$ | 2.45 (1.74–3.44) | $2.21 \times 10^{-7}$ |

†Included all subjects from Chonnam National University (CNU), Samsung Medical Center (SMC), Seoul National University Hospital (SNUH), Sejong University (SU), and Yeonsei University (YSU).

‡Included only females from SNUH, YSU, SU, SMC, and CNU.

§Included subjects from SNU, YSU and SMC.

Abbreviations: CI, confidence interval, OR, odds ratio.

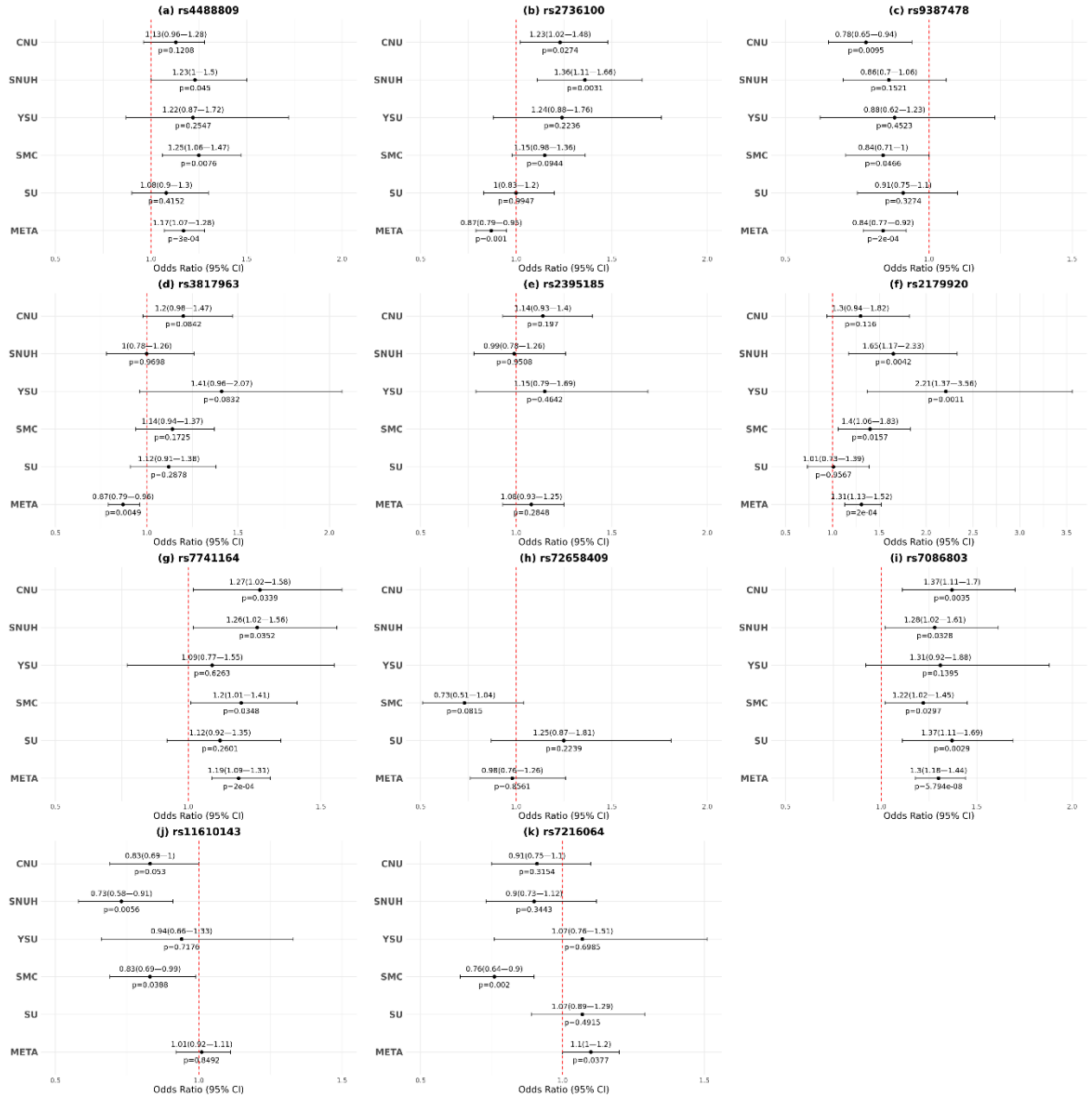


**Figure S1. Estimated odds ratios of the single nucleotide polymorphisms analyzed in each study.** Abbreviations: CI, confidence interval; CNU, Chonnam National University; META, meta-analysis; SMC, Samsung Medical Center; SNUH, Seoul National University Hospital; SU, Sejong University; YSU, Yeonsei University.

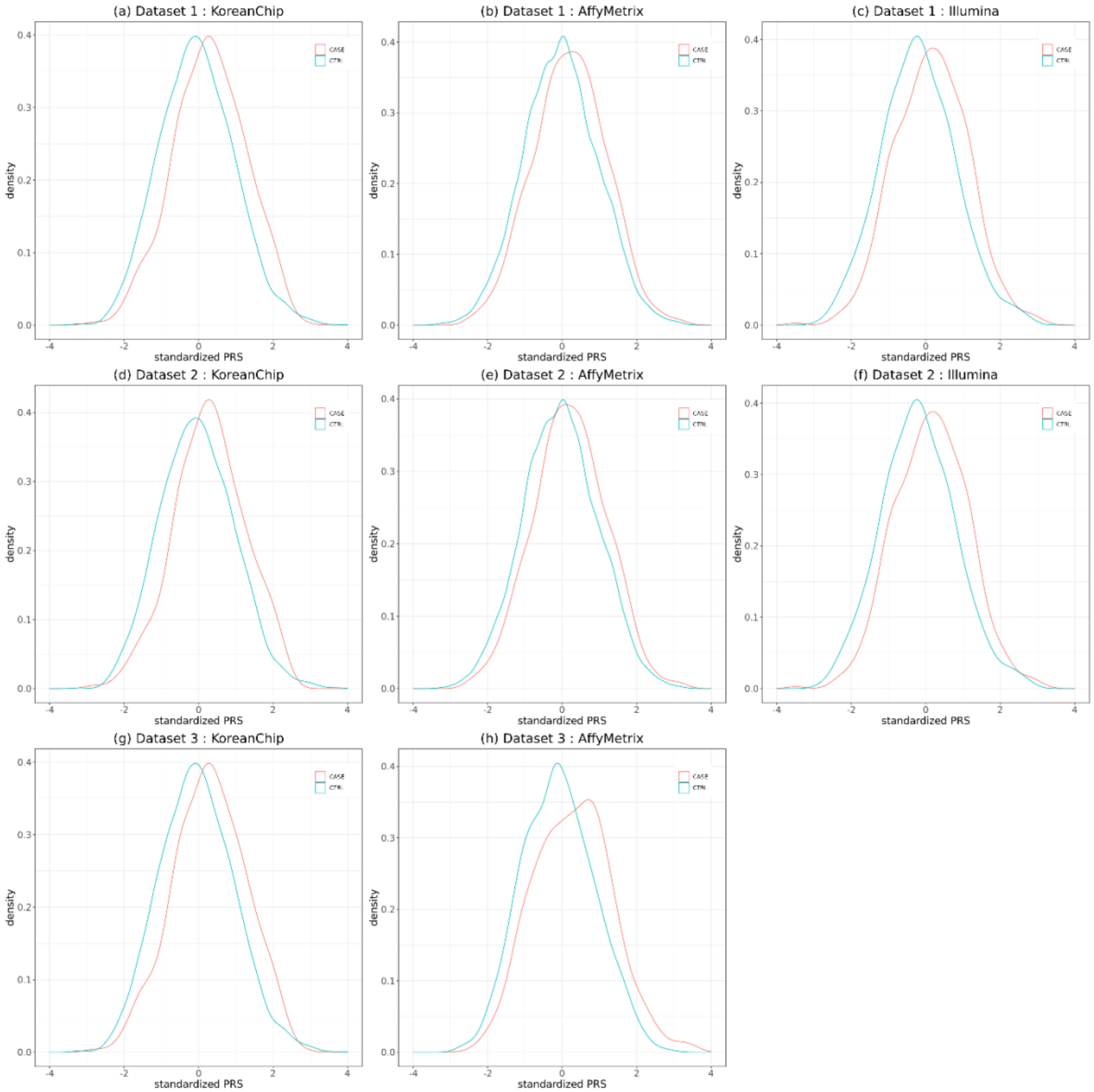


**Figure S2. Distributions of polygenic risk scores (PRSs).** Density plots of standardized PRSs of **(a–c)** Dataset 1, **(d-f)** Dataset 2, and **(g, h)** Dataset 3. Dataset 1 included all subjects from Chonnam National University (CNU), Samsung Medical Center (SMC), Seoul National University Hospital (SNUH), Sejong University (SU), and Yeonsei University (YSU); Dataset 2 included only females from SNUH, YSU, SU, SMC, and CNU; and Dataset 3 included subjects from SNUH, YSU, and SMC.
